# Regulation of the PKD2 Channel Function by TACAN

**DOI:** 10.1101/2021.12.23.473878

**Authors:** Xiong Liu, Rui Zhang, Mohammad Fatehi, Yifang Wang, Wentong Long, Rui Tian, Xiaoling Deng, Ziyi Weng, Qinyi Xu, Peter E Light, Jingfeng Tang, Xing-Zhen Chen

## Abstract

Autosomal dominant polycystic kidney disease (ADPKD) is caused by mutations in membrane receptor PKD1 or cation channel PKD2. TACAN (also named TMEM120A), recently reported as an ion channel in neuron cells for mechano and pain sensing, is also distributed in diverse non-neuronal tissues such as kidney, heart and intestine, suggesting its involvement in other functions. In this study, we found that TACAN is in complex with PKD2 in native renal cell lines. Using the two-electrode voltage clamp in *Xenopus* oocytes we found that TACAN inhibited the channel activity of PKD2 gain-of-function mutant F604P. The first and last transmembrane domains of TACAN were found to interact with the PKD2 C-and N-terminal portions, respectively. We showed that the TACAN N-terminus acted as a blocking peptide and that TACAN inhibits the PKD2 function through the PKD2/TACAN binding. By patch clamping in mammalian cells, we found that TACAN inhibits both the single channel conductance and open probability of PKD2 and mutant F604P. PKD2 co-expressed with TACAN, but not PKD2 alone, exhibited pressure sensitivity. Furthermore, we also found that TACAN aggravates PKD2-dependent tail curvature and pronephric cysts in larval zebrafish, in support of the *in vitro* inhibitory effects of TACAN. In summary, this study revealed that TACAN acts as a PKD2 inhibitor and mediates mechano sensitivity of the PKD2/TACAN channel complex.

## Introduction

Autosomal dominant polycystic kidney disease (ADPKD), characterized by accumulation of multiple cysts in both kidneys, is one of the most common human genetic diseases (Harris and Torres, 2009). Extrarenal pathologies associated with ADPKD may include hepatic and pancreatic cysts, cerebral and intracranial aneurysms, and cardiovascular abnormalities (Luciano and Dahl, 2014). Although mutations in PKD1 or PKD2 or their dosage alterations account for ADPKD, their biophysical and physiological functions are not well understood (Douguet et al., 2019). PKD1 (also called polycystin-1) is a receptor-like membrane protein with 11 transmembrane (TM) segments (S1-S11) and a large extracellular N-terminus while PKD2 (also called polycystin-2 or transient receptor potential polycystin-2 (TRPP2)) is a Ca^2+^-permeable cation channel belonging to the TRP superfamily of cation channels possessing six TMs (S1-S6) and pore domain S5-loop-S6 (Bergmann et al., 2018).PKD2 forms homotetramers but can also for heterotetramers to fulfill different functions (Cheng et al., 2010). As revealed by cryo-electron microscopy (EM) PKD1/PKD2 formed heterotetramers at 1:3 stoichiometry (Su et al., 2018). Through functional studies, we found that PKD1/PKD2 has higher Ca^2+^ permeability than homomeric PKD2 (Wang et al., 2019) but mechanisms of how PKD1 contributes to the selectivity filter and pore gate in PKD1/PKD2 remains elusive. It was reported that the extracellular N-terminus of PKD1 functions as an activation ligand of the PKD1/PKD2 complex (Ha et al., 2020). Additional reports demonstrated that PKD2 interacts with other ion channels including TRPV4 (Köttgen et al., 2008), TRPC1 (Kobori et al., 2009) and Piezo1 (Peyronnet et al., 2013) to form channel complexes with distinct biophysical properties. How PKD2 is regulated by interacting partners remains largely unknown and deserves further studies.

TMEM120A (transmembrane protein 120A, also called TACAN) was initially reported as a nuclear envelope transmembrane protein critical for adipocyte differentiation (Malik et al., 2010). It was recently shown to form a mechano-sensitive ion channel sensing the pain (Beaulieu-Laroche et al., 2020), but this was challenged by subsequent structural and functional studies (Del Rosario et al., 2021; Ke et al., 2021; Niu et al., 2021; Parpaite et al., 2021; Rong et al., 2021; Xue et al., 2021). Global knockout of TACAN led to embryo death indicating its importance for embryonic development (Beaulieu-Laroche *et al.*, 2020). Wide distribution of TACAN in non-neuronal tissues such as kidney, heart and intestine indicate its biological functions besides pain sensation (Beaulieu-Laroche et al., 2020). Previous proteomic screen suggested a potential interaction of PKD2 with TACAN (Sharif-Naeini et al., 2009) but their interaction has yet to be characterized and functional implications to be explored.

In the present study, we examined how TACAN modulates the PKD2 channel function using the two-electrode voltage clamp (TEVC) electrophysiology in *Xenopus* oocytes and single-channel patch clamp electrophysiology in Chinese hamster ovary (CHO) cells. We explored the subcellular localization of endogenous PKD2 and TACAN in different renal cell lines. We also characterized the interaction between TACAN and PKD2 by means of co-immunoprecipitation (co-IP) and immunofluorescence, and explored the relationship between physical association and functional regulation. Further, we examined how TACAN regulates PKD2 deficiency-associated disorders in zebrafish models by clustered regularly interspaced short palindromic repeats/CRISPR-associated protein 9 (CRISPR/Cas9).

## Results

### Effects of TACAN on the PKD2 channel function

TACAN was indicated as a potential interactor of PKD2 through a proteomic screen using smooth muscle cells (Sharif-Naeini et al., 2009). To determine whether and how TACAN modulates the PKD2 channel function, we utilized *Xenopus* oocyte expression together with the two-electrode voltage clamp (TEVC). The whole-cell channel activity of wild-type (WT) PKD2 in oocytes is hardly detectable in part due to its low expression on the plasma membrane and unknown agonist. We instead recorded the current mediated by PKD2 gain-of-function (GOF) mutant F604P (Arif Pavel et al., 2016) in oocytes, similarly as we did previously (Zheng et al., 2018a; Zheng et al., 2018b). We found that oocytes expressing human TACAN alone does not exhibit any significant increase in the current compared with water-injected oocytes (Fig. 1A and B). Co-expression of human PKD2 mutant F604P with TACAN is associated with significantly decreased current amplitudes compared with those with F604P expressed alone (Fig. 1A and B). Because expression of TACAN did not significantly affect the surface expression of F604P, as revealed by whole-cell immunofluorescence (Fig. 1C) and biotinylation assays (Fig. 1D), our data indicated that TACAN inhibits the F604P function. Interestingly, expression of F604P reduced both the total and surface expression of TACAN (Fig. 1E), but the underlying mechanism remains unknown.

**Figure 1.**
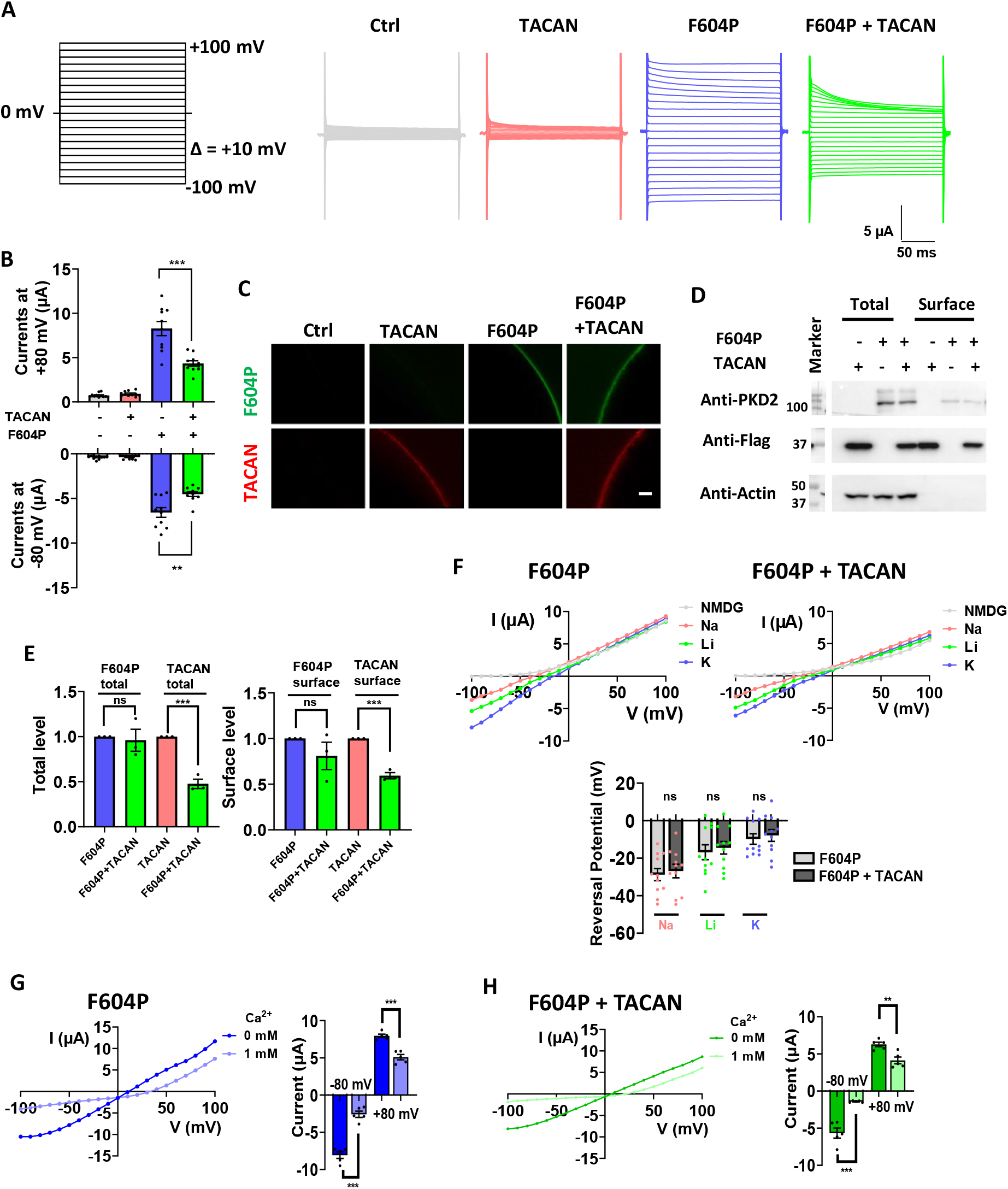
Functional regulation of PKD2 by TACAN in *Xenopus* oocytes. **A.** Representative current traces obtained using a voltage jump protocol in oocytes expressing TACAN, PKD2 F604P (F604P for short) or F604P + TACAN, in the presence of the divalent-free Na-containing solution (in mM): 100 NaCl, 2 KCl, 10 HEPES and pH 7.5. Data from water-injected oocytes served as negative control (Ctrl). **B.** Statistical bar graphs of currents recoded at +80 mV obtained under the same experimental conditions as in **A**. Data are presented as mean ± SEM. N = 10-11 oocytes. * p < 0.05; ** p < 0.01. **C.** Representative IF images showing oocyte surface expression of TACAN or F604P. Scale bar, 50 mm. **D.** Representative Western blot data of the biotinylated (surface) and total protein of F604P and TACAN. **E.** Averaged data obtained as those from **D** (N = 3). Data are presented as mean ± SEM. ns, not significant; *** p < 0.001. **F.** Upper panels: representative I-V curves for F604P or F604P + TACAN when 100 mM of an indicated cation was used in the divalent ion-free bath solution. Lower panel: averaged reversal potentials. Data are presented as mean ± SEM. N = 11 oocytes. ns, not significant. **G** and **H.** Left panel: representative I-V curves for PKD2 F604P (**G**) and F604P + TACAN (**H**) in bath solution with or without 1 mM Ca^2+^. Right panel: effect of Ca^2+^ on the inward (−80 mV) and outward (+80 mV) currents for F604P (**G**) and F604P + TACAN (**H**). Data are presented as mean ± SEM. N = 5 oocytes. * p < 0.05; ** p < 0.01; *** p < 0.001.

Next, we examined whether TACAN affected the F604P ion selectivity using extracellular solutions containing 100 mM Na^+^, Li^+^, K^+^ or non-permeable N-methyl-d-glucamine (NMDG, negative control). We found that the reversal potentials obtained under different solutions for F604P + TACAN were not significantly different from those for F604P (Fig. 1F), indicating that TACAN does not significantly affect the cation selectivity (with permeability ratios P_Na_: P_Li_: P_K_ = 1: 1.65: 2.15 for F604P, consistent with previous reports (Shen et al., 2016), and P_Na_: P_Li_: P_K_ = 1: 1.66: 2.15 for F604P + TACAN). Extracellular Ca^2+^ is known to reduce the PKD2 conductance to monovalent cations (Arif Pavel et al., 2016; Shen et al., 2016). Here we found that Ca^2+^ exhibits a similar inhibitory effect on F604P + TACAN and F604P alone (Fig. 1G and H). Taken together, our data showed that TACAN inhibits the mutant F604P channel activity but does not affect the cation selectivity or inhibition of F604P by extracellular Ca^2+^.

To further characterize how TACAN inhibits the PKD2 channel function, we performed single-channel patch clamp electrophysiology in CHO cells. Expression of TACAN with mutant F604P or WT PKD2 substantially reduced the single-channel amplitude and open probability while TACAN expressed alone did not induce any specific current under our experimental condition (Fig. 2A and B), consistent with whole-cell data obtained using oocytes. We also noticed that at +80 mV in the absence of TACAN the single-channel amplitude and open probability values for PKD2 are significantly lower than those for mutant F604P, which together with a lower density of PKD2 on the surface membrane may account for its much lower whole-cell currents.

**Figure 2.**
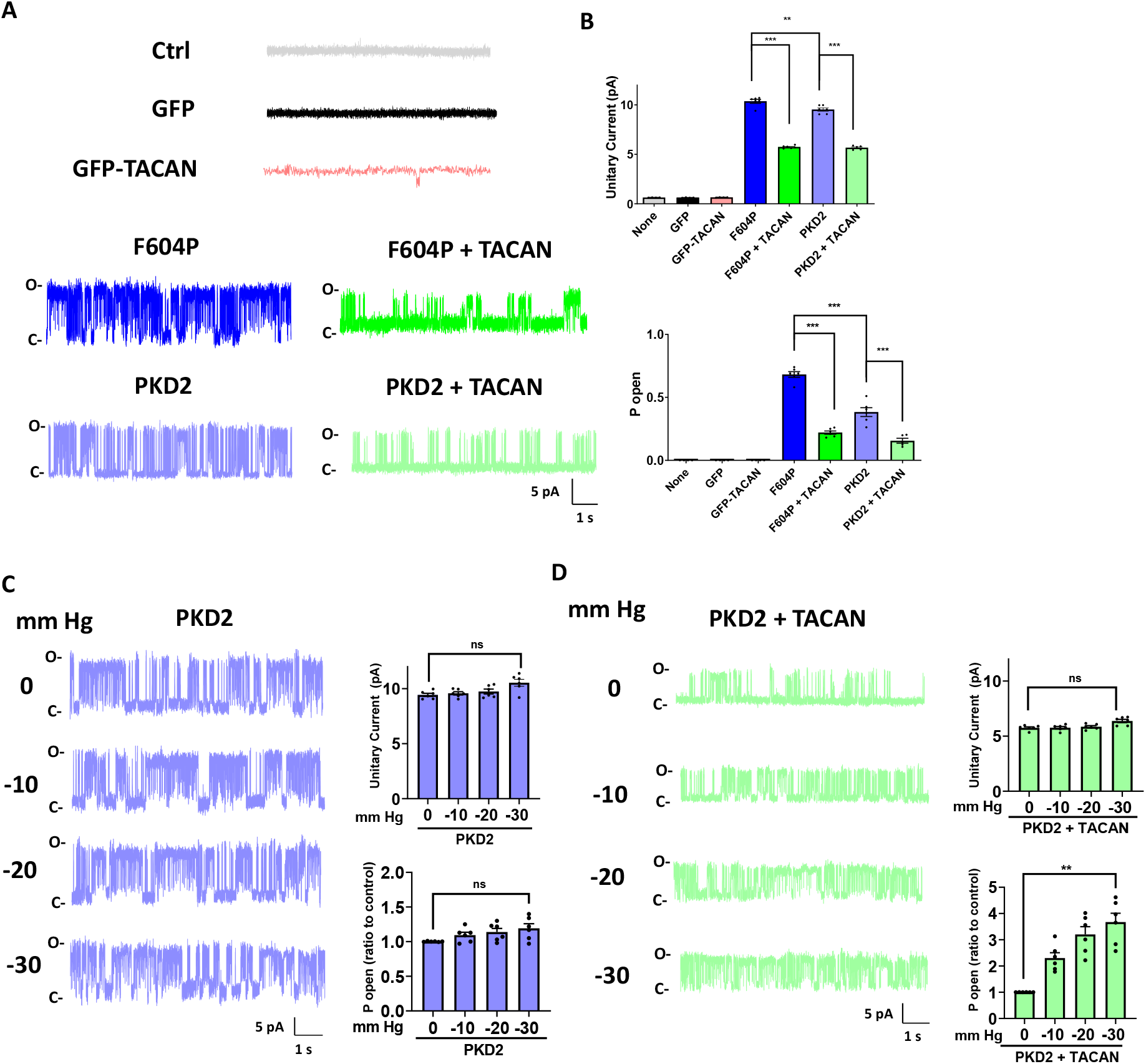
Effects of TACAN and pressure on PKD2 single-channel parameters in CHO cells. **A.** Representative single-channel recordings (at +80 mV) of CHO cells with indicated transfections. The same solution (in mM: 140 NaCl, 5 KCl, 2 CaCl_2_, 2 MgCl_2_, 10 HEPES, 10 glucose, and pH 7.4) was used in the pipette and bath. **B.** Quantified unitary current amplitudes and open probabilities obtained from **A**. Data are presented as mean ± SEM. N = 4-6 cells. ** p < 0.01; *** p < 0.001. **C** and **D.** Left panels: representative single-channel recordings (at +80 mV) in CHO cells transfected with PKD2 (**C**) or PKD2 + TACAN (**D**) showing the effect of negative pressures. Right panels: effect of the pressure on the unitary currents and normalized open probabilities of PKD2 (**C**) or PKD2 + TACAN (**D**). Data are presented as mean ± SEM. N = 6 cells. ns, not significant; ** p < 0.01.

We next examined pressure sensitivity of PKD2 in CHO cells with or without co-expression of TACAN. PKD2 in complex with PKD1 or TRPV4 was reported to be involved in mechano sensing (Köttgen et al., 2008; Nauli et al., 2003) but whether PKD2, PKD1, TRPV4 or an unknown binding protein senses mechanical stimuli is not well understood. With PKD2 expressed alone in CHO cells, pressure up to −30 mm Hg had no effect on the single-channel amplitude or open probability (Fig. 2C). In contrast, in the presence of TACAN co-expression, the open probability value significantly increased with negative pressure while the single-channel current amplitude was insensitive to the pressure (Fig. 2D). These data showed that channel complex PKD2/TACAN possesses mechano sensitivity mediated by TACAN, which the single-channel open probability, but not the unitary current, stimulated by the pressure. Of note, no channel activity was observed in cells expressing TACAN alone with pressure up to −30 mm Hg (Fig. S1). Increasing the pressure beyond −30 mm Hg in our setup generated non-specific leak currents, presumably due to broken cell membrane.

Besides inhibiting the F604P steady-state currents TACAN induced significant inactivation of the F604P-mediated currents at depolarization (Fig. 3A). The steady-state current to the peak current ratio for F604P alone was close to 1 (1.01 ± 0.01 at +100 mV, N = 7, p = 0.73), ie, there was no appreciable inactivation, while this ratio decreased to 0.67 ± 0.04 (N = 7, p < 0.001) with TACAN co-expression (Fig. 3B). The current inactivation at depolarizations (+50 to +100 mV) in the presence of TACAN was voltage dependent, with time constant (τ) values in the range of 139.23 ± 15.43 and 33.8 ± 4.80 ms (N = 7) obtained from exponential fits (Fig. 3C). Interestingly, significant inactivation at polarized voltages was also present in PKD2 GOF gate mutant L677G expressed alone (Fig. 3D). Unlike the F604P mutation, which locks PKD2 in an activated configuration, the GOF of the L677G mutation is due to direct increases in the pore hydrophilicity and size, which would infer that the L677G mutant protein may still be in a basal configuration similar to WT PKD2 (Zheng et al., 2018b). Interestingly, although TACAN significantly reduced the steady-state currents of L677G (Fig. 3D–F, it did not affect the inactivating part of the L677G currents, which were associated with a similar range of inactivation time constant compared with F604P + TACAN (Fig. 3G vs Fig. 3C). These data together seemed to suggest that inactivation occurs when PKD2 protein is in an inhibited (F604P + TACAN) or a basal state (L677G, with or without TACAN) but not in an activated state (F604P alone).

**Figure 3.**
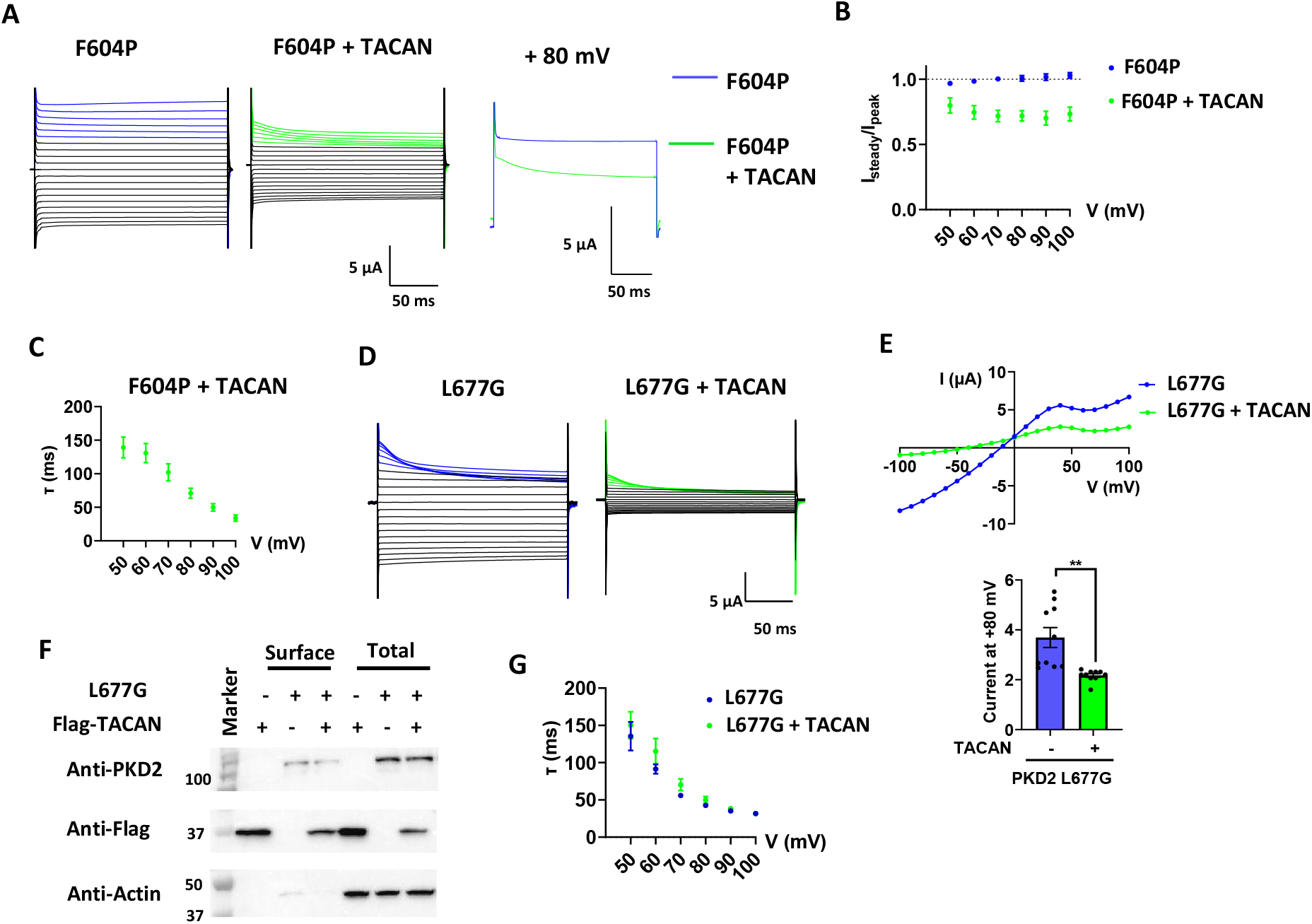
Effect of TACAN on inactivation of PKD2-mediated currents in oocytes. **A.** Representative traces (from three independent experiments) from oocytes expressing PKD2 F604P or F604P + TACAN. Traces obtained from +50 mV to +100 mV were marked blue or green, as indicated. For clarity, those at +80 mV were compared. **B.** Ratios of the steady currents (I_steady_) to the peak currents (I_peak_) for F604P and F604P + TACAN obtained as in **A** (N = 7 oocytes). **C.** Time constant (τ) of current decay for F604P + TACAN obtained through an exponential fit (N = 7 oocytes). **D.** Representative traces (from three independent experiments) from oocytes expressing PKD2 L677G or L677G + TACAN. Traces obtained from +50 mV to +100 mV are marked blue or green, as indicated. **E.** Left panel: Representative I-V curves from oocytes expressing L677G or L677G + TACAN. Right panel: averaged currents at +80 mV obtained from oocytes expressing L677G or L677G + TACAN. Data are presented as mean ± SEM. N = 10 oocytes. ** p< 0.01. **F.** Representative Western blot data of the surface (biotinylated) and total proteins of L677G and TACAN. **G.** Time constants (τ) of current decay for L677G and L677G + TACAN (N = 5 oocytes).

### Physical interaction between PKD2 and TACAN

TACAN was shown to be highly expressed in the kidney (Beaulieu-Laroche et al., 2020). We wanted to check the subcellular localization of PKD2 and TACAN be means of double immunofluorescence assays in ciliated epithelia Madin-Darby Canine Kidney (MDCK), inner medullary collecting duct (IMCD), and Lilly Laboratories Cell-Porcine Kidney 1 (LLC-PK1) cell lines. We found that the endogenous PKD2 and TACAN both display enhanced staining in the primary cilia in all the three cell lines (Fig. 4A), indicating co-distribution of the two proteins. We also performed co-IP experiments using these cell lines to assess their interaction and found that PKD2 is present in the immunoprecipitates obtained with an anti-TMEM120A antibody, but not in those with IgG (control) (Fig. 4B), demonstrating that PKD2 and TACAN are in the same protein complex in these native cells.

**Figure 4.**
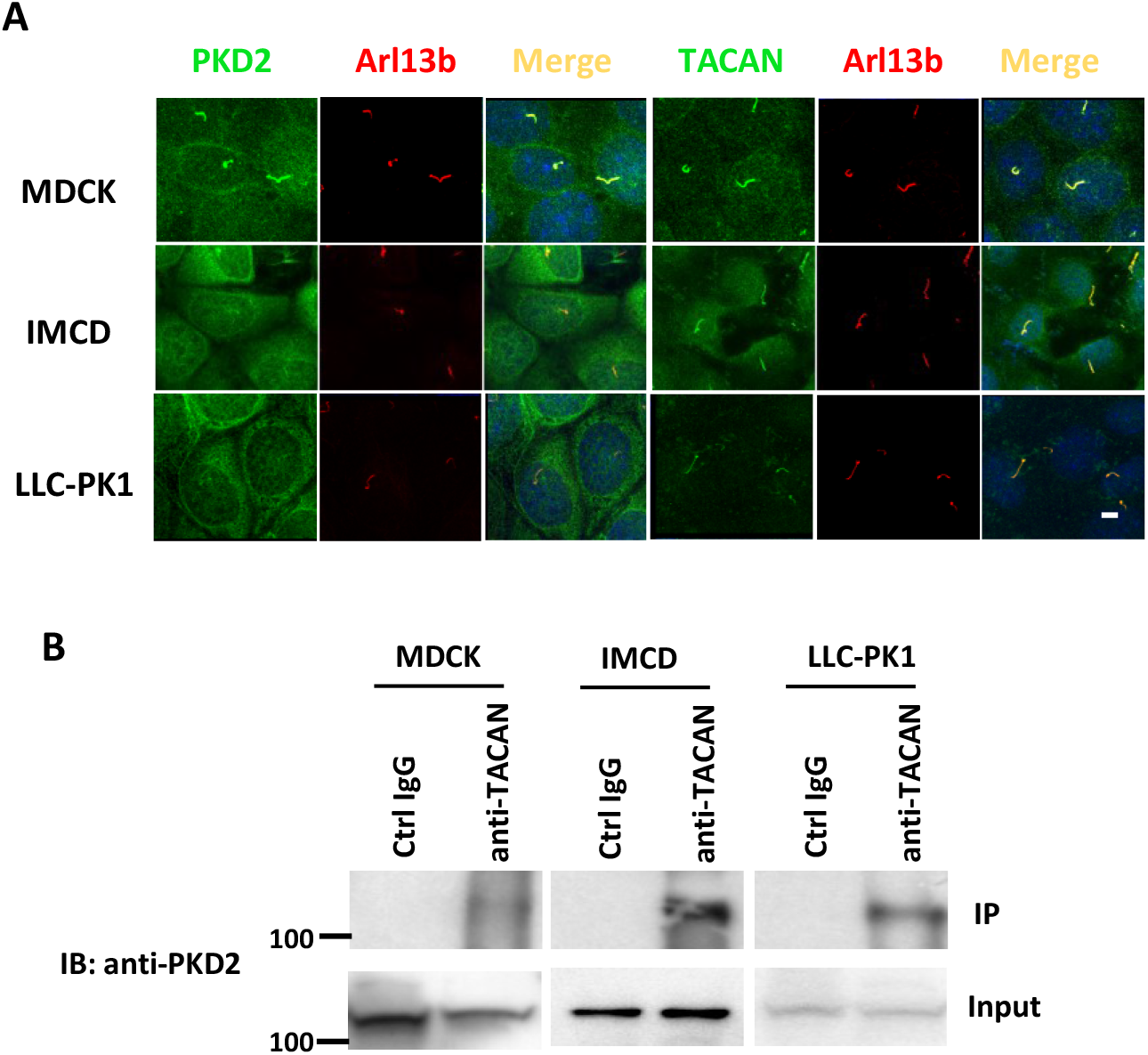
Colocalization and association of the endogenous PKD2 and TACAN in renal cell lines. **A.** Subcellular localization of the endogenous PKD2 and TACAN in MDCK, IMCD and LLC-PK1 cells by means of immunofluorescence. Arl13b served as a primary cilia marker. Scale bar, 5 μm. Shown are representative data from three independent experiments. **B.** Representative co-IP data (from three independent experiments) showing association between the endogenous PKD2 and TACAN in MDCK, IMCD and LLC-PK1 cells.

We next wanted to characterize the physical interaction between PKD2 and TACAN interaction in oocytes to better relate to our functional data. Indeed, TACAN co-immunoprecipitated TACAN and mutant F604P in oocytes and reversely, mutant F604P was able to precipitate TACAN (Fig. 5A–C), indicating that PKD2 and TACAN are in the same complex in oocytes. We then used chemical cross-linking to examine the subunit stoichiometry of the PKD2/TACAN complex. However, the complex cannot be detected under our conditions although we successfully detected TACAN dimers and different PKD2 oligomers (Fig. S2A). The coiled-coil domain (amino-acid (aa) G9-L100) in the TACAN N-terminus was proposed to be important for oligomerization (Batrakou et al., 2015). Indeed, we found aggregation of the N-terminus in our chemical cross-linking experiments. Further, TACAN lacking the entire N-terminus still formed the dimer (Fig. S2B), indicating that the TACAN dimeric assembly is independent of its coiled-coil domain. Furthermore, we generated different fragments of TACAN, which possesses six TMs (S1-S6), to narrow down the domain(s) mediating physical and functional interaction with PKD2. By co-IP assays we found that TACAN fragments containing S1 (K135-T160) or S6 (L296-D343), but not the N-terminus (K135X), interacts with PKD2 F604P (Fig. 5D). While S1-containing TACAN fragments exhibited no functional effect on F604P, interestingly, S6-containing fragments lacking the N-terminus exhibited significant inhibitory effects (Fig. 5E). Further, this inhibition was even stronger than the one by full-length TACAN (Fig. 5E), presumably because these fragments can’t form dimers (Fig. S2B), which enhanced their interaction with F604P.

**Figure 5.**
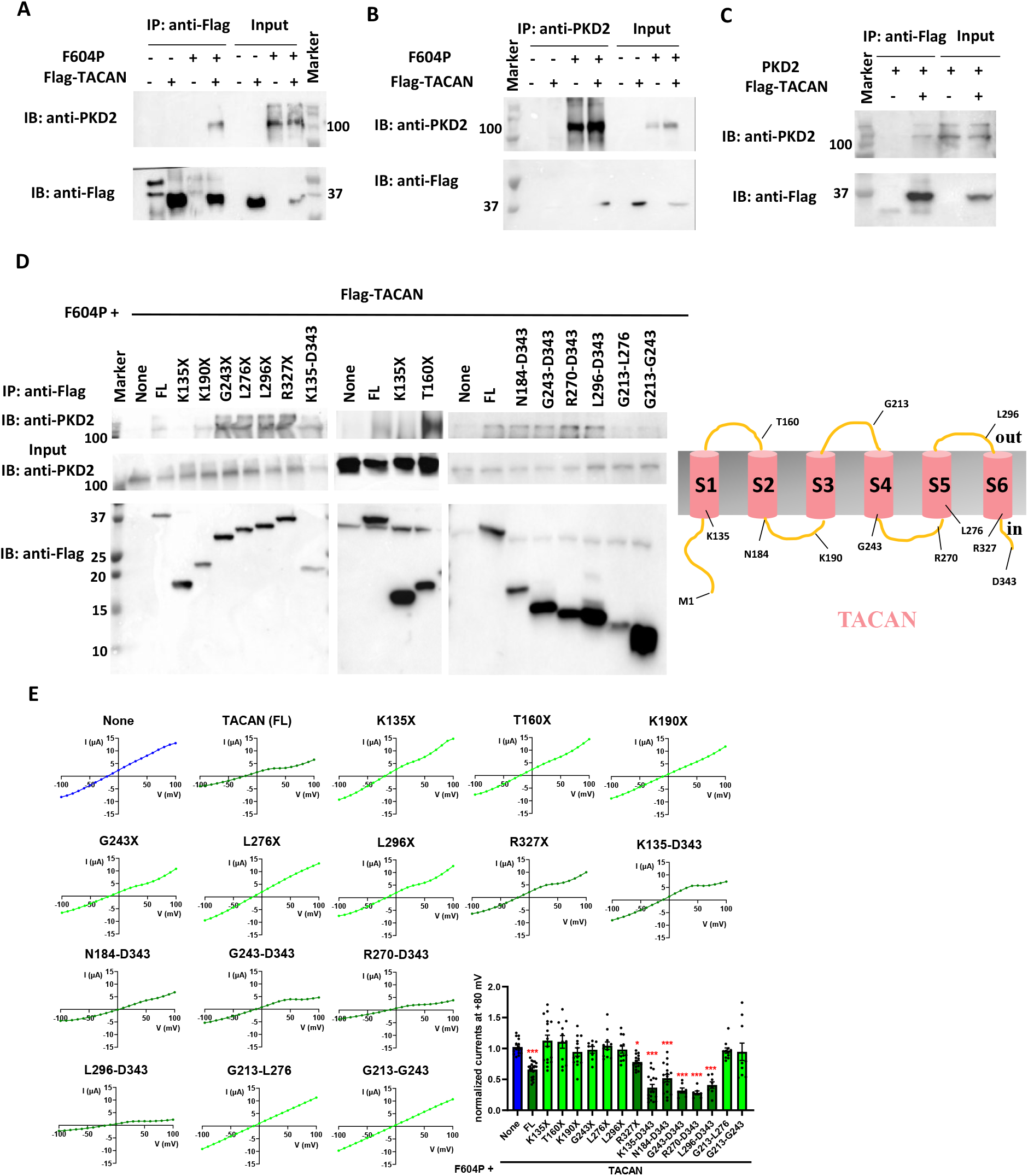
Domains of TACAN mediating association with PKD2. **A** and **B.** Representative co-IP data (from three independent experiments) showing the interaction between PKD2 mutant HA-F604P and Flag-TACAN using Flag antibody (**A**) or PKD2 antibody (**B**) for precipitation. **C.** Representative co-IP data (from three independent experiments) showing interaction between WT HA-PKD2 and Flag-TACAN using Flag antibody for precipitation. **D.** Left panels: representative co-IP data (from three independent experiments) showing the interaction between mutant HA-F604P and an indicated TACAN truncation mutant using Flag antibody for precipitation. Right panel: topology of human TACAN. Indicated residue numbers stand for points of truncation mutations. **E.** Representative I-V curves and averaged currents at +80 mV in oocytes expressing F604P with or without a TACAN truncation mutant. Data are presented as mean ± SEM. N = 7-20 oocytes. * p < 0.05; ** p < 0.01; *** p < 0.001.

Reversely, we wanted to identify which part(s) in PKD2 is involved in the interaction with TACAN. We performed co-IP experiments in CHO cells expressing TACAN together with the PKD2 N-terminus (PKD2-N, M1-K215), C-terminus (PKD2-C, D682-V968), or PKD2 lacking both the N- and C-termini (PKD2-TM, S209-K688). PKD2-TM but not PKD2-N or PKD2-C conferred interaction with TACAN (Fig. 6A), which challenges the previous proteomic screen study indicating that TACAN binds to the PKD2 C-terminus (Sharif-Naeini et al., 2009).

**Figure 6.**
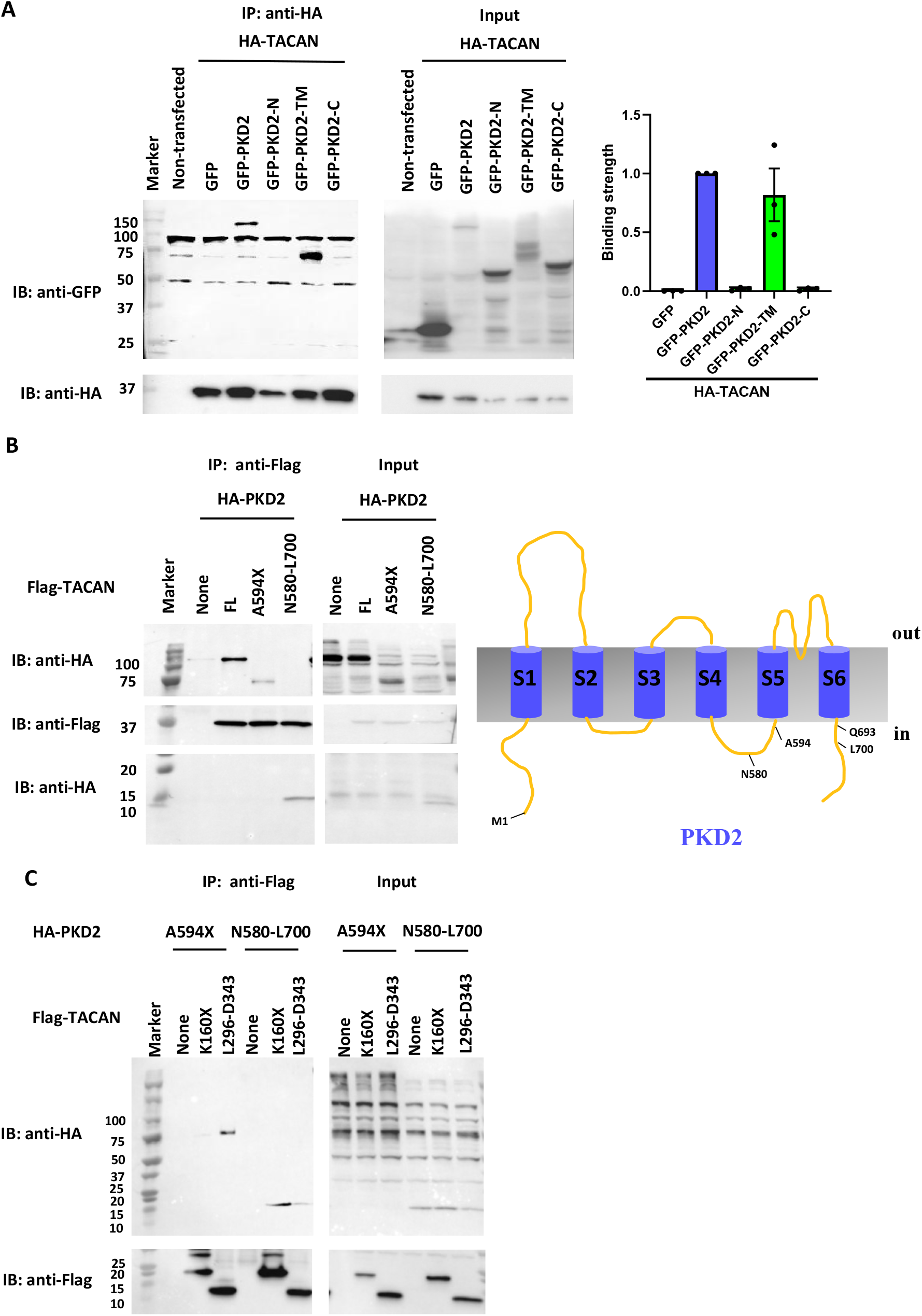
Domains in PKD2 mediating association with TACAN. **A.** Left panels: representative co-IP data (from three independent experiments) showing the interaction between GFP-PKD2 and HA-TACAN using HA antibody for precipitation. Non-transfected and GFP-transfected CHO cell lysates served as negative controls. Right panel: binding strength averaged from three independent experiments. **B.** Left panels: representative co-IP data (from three independent experiments) showing the interaction between Flag-TACAN and a PKD2 truncation mutant (A594X or N580-L700) using Flag antibody for precipitation. PKD2 expressed in oocytes with or without co-expressed Flag-TACAN served as positive and negative controls, respectively. Right panel: topology of human PKD2. Indicated residue numbers stand for points of truncation mutations. **C.** Representative co-IP data (from three independent experiments) showing the interaction between PKD2 mutant A594X and TACAN mutant L296-D343, and between PKD2 mutant N580-L700 and TACAN mutant K160X using Flag antibody for precipitation. PKD2 mutants A594X and N580-L700 alone served as negative controls.

The PKD2 pore domain is formed by S5, S5-S6 loop and S6 (A594-Q693) (Shen et al., 2016). We next examined whether TACAN participates in the pore formation in the PKD2/TACAN channel complex. For this purpose, we tested whether TACAN interacts with truncation mutant A594X (no pore domain) or fragment N580-L700 containing the pore domain. Surprisingly, TACAN can interact with either of them (Fig. 6B), indicating that PKD2 also contains at least two domains interacting with TACAN. Because the TACAN S1 and S6 interacted with PKD2, we continued to narrow down the corresponding binding domains in PKD2. We found by co-IP that PKD2 A594X interacts with TACAN S6, while PKD2 N580-L700 interacts with TACAN S1 (Fig. 6C). Since the interaction between S6 was functionally critical, it highly suggested that instead of acting as the pore formation subunit, TACAN mainly binds at the peripheral domain (S1-S4) to regulate PKD2 channel function.

We wondered whether the PKD2/TACAN binding is required for inhibition of the PKD2 function by TACAN. For this, we co-expressed the TACAN N-terminus (K135X) in the presence of F604P and TACAN, and indeed found that K135X represses the PKD2/TACAN association (Fig. 7A), indicating that K135X acts as an effective blocking peptide, presumably because it can compete with PKD2 for binding (dimerizing) with TACAN (see Fig. S2). Further, K135K which had no functional effect on PKD2 (Fig. 5E) significantly represses the inhibition of the PKD2 function by TACAN (Fig. 7B). Our data together thus demonstrated that inhibition of the PKD2 function by TACAN is mediated by their physical association.

**Figure 7.**
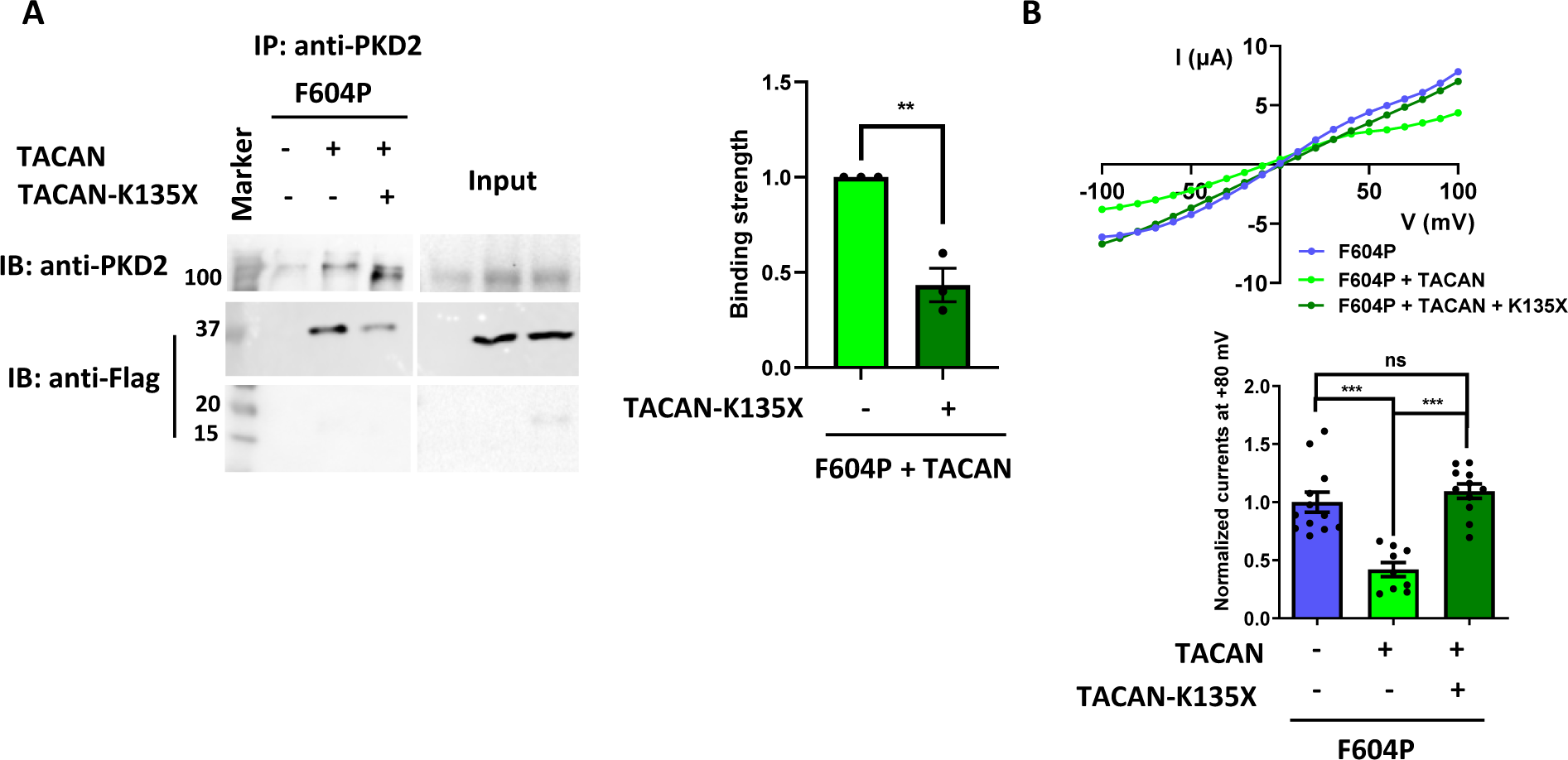
Effect of the TACAN N-terminus on TACAN’s inhibition of the PKD2 function. **A.** Left panel: representative co-IP data showing the interaction between F604P and Flag-TACAN, with or without TACAN N-terminus (K135X) in oocytes, using PKD2 antibody for precipitation. Right panel: binding strength averaged from three independent experiments. **B.** Upper panel: representative I-V curves from oocytes expressing F604P, F604P + TACAN, or F604P + TACAN + K135X. Lower panel: averaged currents at +80 mV. Data are presented as mean ± SEM. N = 9-12 oocytes. ns, not significant, *** p < 0.001.

### Regulation of the PKD2 function in zebrafish by TACAN

We next utilized zebrafish models to investigate the regulatory role *in vivo* of TACAN in PKD2. Sufficient reduction in the PKD2 expression or function in larval zebrafish results in the presence of tail curvature and pronephric cysts resembling renal cysts in mammals (Zheng et al., 2016). Here, we employed the clustered regularly interspaced short palindromic repeats/CRISPR-associated protein 9 (CRISPR/Cas9) technique and successfully knocked down the endogenous PKD2 in larval fish at 3 days post-fertilization (dpf) through injection of 200 pg PKD2 sgRNA and 300 pg Cas9 per embryo, which resulted in tail curling (Fig. 8A and B). We reasoned that if we reduce the injected PKD2 sgRNA and Cas9 amounts there may be no or low occurrence of tail curling but co-expression of TACAN may significantly aggravate the severity of the disease if the PKD2 function is inhibited.

**Figure 8.**
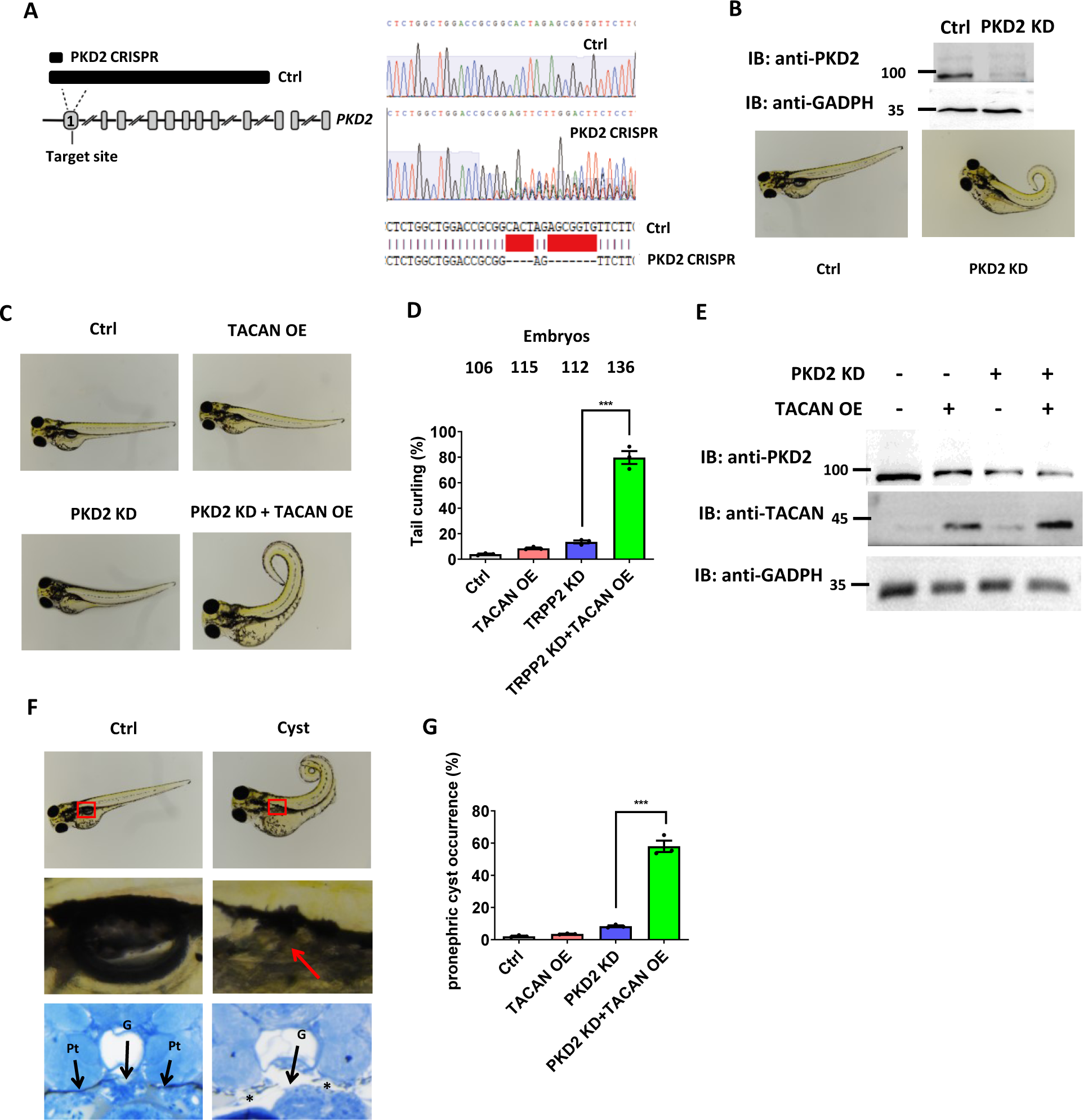
Effect of TACAN on the tail curling and pronephric cystogenesis of larval zebrafish. **A.** Left panel: schematic presentation of the PKD2 protein/gene domain architecture and CRISPR target sites. The gene loci are shown with exons (gray boxes) and introns (solid lines). The position of the CRISPR target sites in exon 1 in the PKD2 gene and the corresponding PKD2 protein (black box) are indicated by the dash lines, and the predicted remaining (truncated) PKD2 protein (PKD2 CRISPR) is indicated. Right panel: sequencing of PKD2 in the Ctrl and PKD2 CRISPR fish at 3 dpf. Red lines indicate CRISPR target sites. **B.** Upper panel: PKD2 protein detected by Western blot under the Ctrl (water injection) and CRISPR-directed PKD2 knockdown (KD) conditions (PKD2 sgRNA 200 pg + Cas9 300 pg) at 3 dpf. Lower panels: representative pictures of embryos at 3 dpf. **C.** Representative pictures showing tails of 3 dpf zebrafish under the Ctrl, TACAN overexpression (OE) (by injection of mRNA at100 pg each), or PKD2 KD condition (PKD2 sgRNA 100 pg + Cas9 200 pg). **D.** Average percentages of fish displaying curled tail under different conditions. Data were from three independent experiments with the indicated total numbers of embryos. ns, not significant, *** p < 0.001. **E.** Representative western blot data (from three independent experiments) showing the expression of PKD2 and TACAN. **F.** Representative pictures of a 3-dpf embryo with PKD2 KD plus TACAN OE, showing pronephric cyst formation (red box and arrow). Ctrl fish was injected with water only. Cyst formation was also indicated by a histologic section showing dilated pronephric tubules (asterisks). G, glomerulus; Pt, pronephric tubule. **G.** Averaged percentages of embryos exhibiting pronephric cysts under the indicated conditions. Data are presented as mean ± SEM. *** p < 0.001.

Indeed, while injection of 100 pg PKD2 sgRNA and 200 pg Cas9 per embryo was associated with low percentage of fish with tail curling, co-expression of human TACAN through co-injection on *in vitro* transcribed mRNA (100 pg/embryo) substantially increased the occurrence of tail curling (Fig. 8C and D), consistent with the assumption that the PKD2 function is inhibited by over-expressed TACAN, which was observed in oocytes and CHO cells. Our Western blot experiments showed that the PKD2 expression is not significantly reduced by TACAN over-expression (Fig. 8E), which is in agreement with our data from cellular models. We also observed that TACAN over-expression alone did not significantly increase the occurrence of tail curling (Fig. 8D), indicating that the functional inhibition of the endogenous PKD2 by TACAN is in general insufficient to induce the disease. We also examined pronephric cysts in larval fish and found that TACAN co-expression substantially increases the occurrence of pronephric cysts while TACAN over-expression or PKD2 knockdown alone exhibits no or low occurrence of pronephric cyst (Fig. 8F and G). Taken together, our data demonstrated that TACAN aggravates PKD2-dependent disease severity in larval zebrafish, presumably through repressing the PKD2 channel function.

## Discussion

TACAN is a transmembrane protein that is evolutionally conserved and was recently reported to be involved in sensing mechanical stimuli (Beaulieu-Laroche et al., 2020). Its intracellular N-terminal coiled-coil domain (aa G9-L100) was reported to be important for oligomerization (Batrakou et al., 2015), consistent with our cross-linking results showing that deletion of the N-terminus significantly reduced its dimerization. However, the remaining dimers indicate the presence of another domain important for dimerization. While this other domain involved in homo-dimerization remains unknown, our current study showed that the TACAN TM domains S1 and S6 mediate binding with PKD2. Interestingly, while the PKD2 C-terminal coiled-coil domain was reported to be involved in interaction with PKD1 (Qian et al., 1997; Tsiokas et al., 1997; Yu et al., 2009), heterotetrameric PKD1/PKD2 structure was revealed by cryo-EM in the absence of the PKD2 coiled-coil domain (Su et al., 2018), demonstrating that the domain is not essential for the PKD1/PKD2 complexing. In fact, we here showed that the PKD2 TMs but not its N- or C-terminus interacts with the TACAN S1 and S6.

Besides homotetramerization, PKD2 can heteromerize with PKD1 and TRP channels including TRPC1, −C3, −C4, −C7 and −V4 (Cheng et al., 2010; Grieben et al., 2017; Shen et al., 2016). PKD2 homo-and heterotetrameric channels have distinct biophysical properties as well as distinct subcellular localizations (Cheng et al., 2010). The PKD1/PKD2 heterotetrameric channel defective in ADPKD has a 1:3 stoichiometry (Su et al., 2018). While it is well known that PKD1 promotes the plasma membrane trafficking of PKD2, how it contributes to the channel pore formation is not well understood (Ha et al., 2020; Wang et al., 2019). PKD1/PKD2 located in renal epithelial primary cilia was previously shown to mediate mechano-dependent Ca^2+^ entry(Nauli et al., 2003) but subsequent studies found that, controversially, the Ca^2+^ entry is PKD1-independent (Liu et al., 2018). PKD2/TRPC4 in the MDCK primary cilia was found to be involved in flow sensing but the role of primary cilia and mammalian TRP channels in mechano sensing remains debatable (Delling et al., 2016; Köttgen et al., 2008; Nikolaev et al., 2019). Our current study found that in the PKD2/TACAN channel complex TACAN but not PKD2 confers mechano sensitivity (Fig. 2D). It will be interesting to investigate whether TACAN plays a role in the pathogenesis of ADPKD.

Mammalian TACAN (TMEM120A) and its homologue TMEM120B (70.5% sequence identity) are generated as a result of gene duplication. However, when they were expressed in CHO cells, only TACAN induced mechano-sensitive currents (Beaulieu-Laroche et al., 2020). TACAN, but not TMEM120B, was reported to inhibit the Piezo2 channel function (Del Rosario et al., 2021). It will be of interest to determine whether TMEM120B regulates the PKD2 function.

Zebrafish has a relatively simple renal system called pronephros that develop within 24 hours post-fertilization (hpf) and represents an established PKD model with easy visualization of organs and tissues with transparent embryos and larvae, and the ability for rapid phenotype analysis. PKD2 knockdown (KD) by injection of morpholino antisense oligonucleotides (MO) in zebrafish embryos results in tail curling and pronephric cysts at 3-5 days post-fertilization (dpf) (Zheng et al., 2016). However, because MO knockdown may have off-target effects, in the present study we knocked down PKD2 by CRISPR/Cas9, which is known to be more specific than MO knockdown. Our study using larval zebrafish demonstrated that TACAN aggravates the severity of PKD2 insufficiency-dependent diseases, presumably through repressing the PKD2 function but not the expression, consistent with its regulatory effect on PKD2 using cellular models.

Interestingly, it was reported that TMEM33, an activator of PKD2, which forms a complex on the endoplasmic reticulum (ER) membrane, but not in primary cilia, fails to regulate PKD2-dependent pronephric renal cystogenesis in zebrafish (Arhatte et al., 2019), indicating that PKD2 located on the ER membrane is not involved in cystogenesis. Thus, the colocalization or interaction of PKD2 and TACAN in the renal primary cilia may be important for TACAN to confer its regulatory role in PKD2 and the associated disease phenotypes in zebrafish.

The report that TACAN acts as an ion channel has been challenged by recent structural and functional studies (Beaulieu-Laroche et al., 2020; Del Rosario et al., 2021; Ke et al., 2021; Niu et al., 2021; Parpaite et al., 2021; Rong et al., 2021; Xue et al., 2021). However, the concept that TACAN is an ion channel is supported by a recent report showing that a point mutation at a candidate pore gate residue significantly increases the channel activity (Chen et al., 2021) and that, by molecular dynamic simulations, each TACAN monomer contains an ion conducting pathway but multiple subunits might be required for TACAN to form a functional ion channel. Also, TACAN was shown to be involved in regulation of ion transport based on its involvement in regulation of the expression of different proteins including Ca^2+^ ATPase, ATP2a1 and TMEM150C, a general regulator of mechano-sensitive channels such as Piezo1, Piezo2 and the potassium channel TREK-1 (Anderson et al., 2018). Structural studies found that TACAN structure resembles that of the long-chain fatty acid elongase 7 (Ke et al., 2021; Niu et al., 2021; Rong et al., 2021; Xue et al., 2021), suggesting the possibility that TACAN may act as an enzyme. In addition to our current finding, TACAN was previously reported to be an inhibitor of Piezo2, but not Piezo1 or TREK-1, but with an unclear mechanism of inhibition (Del Rosario et al., 2021).

In summary, our present study found that TACAN inhibits the PKD2 function through the physical PKD2/TACAN association. It is unlikely that TACAN participates in the pore formation in the PKD2/TACAN channel complex since TACAN-S6 that doesn’t seem to be sufficient for contributing to pore formation still exhibited an inhibitory effect. As a novel regulator of PKD2, TACAN inhibits PKD2 *in vitro* and *in vivo*. Blocking peptides such as K135X may eventually lead to specific peptide molecules with therapeutic potential for ADPKD.

## Methods

### Plasmids, reagents and antibodies

HA-tagged human PKD2 (HA-PKD2) in vector pGEMHE for *Xenopus* oocyte expression and PKD2 mutant F604P in vector pcDNA3.1 for mammalian cell expression were obtained from Dr. Yong Yu (St. John’s University, NY). Human HA-TACAN in a modified pCMV vector for mammalian cell expression, a kind gift from Dr. Reza Sharif-Naeini (McGill University, QC, Canada), was subcloned into vector PGEMHE for oocyte expression, with a 5’ Flag tag. Human PKD2, PKD2-N, PKD2-TM and PKD2-C in vector pEGFP for oocyte expression were constructed previously (Wang et al., 2012). All mutations were carried out using Q5 High-Fidelity 2X Master Mix from New England Biolabs (NEB, Ottawa, ON, Canada) and verified by sequencing. Antibodies against β-actin and Arl13b were from Santa Cruz Biotechnology (Santa Cruz, CA) and those against Flag, HA, TMEM120A and PKD2 were from Proteintech Group (Rosemont, IL). Secondary antibodies were purchased from GE Healthcare (Waukesha, WI). DTT was purchased from Thermo Fisher Scientific (Ottawa, ON, Canada) and disuccinimidyl tartrate (DST) from CovaChem (Loves Park, IL).

### *Xenopus* oocyte expression

Capped RNAs encoding human PKD2, human TACAN or their mutants were synthesized by *in vitro* transcription using the T7 mMESSAGE mMACHINE kit (Invitrogen, Waltham, MA) and injected into oocytes (25 ng each), as described (Cai et al., 2020). Control oocytes were injected with the same amount of water. Oocytes were incubated at 18 °C for 2-3 days before measurements. The present study was approved by the Ethical Committee for Animal Experiments of the University of Alberta and performed in accordance with the Guidelines for Research with Experimental Animals of the University of Alberta and the Guide for the Care and Use of Laboratory Animals (NIH Guide) revised in 1996.

### Culture and transfection of mammalian cells

CHO cells were cultured in Dulbecco’s Modified Eagle Medium/Nutrient Mixture F-12 (DMEM/F12) supplemented with L-glutamine, penicillin-streptomycin and 10% fetal bovine serum (FBS). MDCK, IMCD and LLC-PK1 cells were cultured in DMEM supplemented with L-glutamine, penicillin-streptomycin and 10% FBS. All cultured cells were kept at 37°C with 5% CO_2_. Transfection of cDNAs was performed using Lipofectamine 3000 (Invitrogen) according to the manufacturer’s protocol.

### Two-electrode voltage clamp

The two-electrode voltage clamp electrophysiology experiments in *Xenopus* oocytes were performed as we described previously (Zheng et al., 2018a). Briefly, an electrode made of a capillary glass pipette (Warner Instruments, Hamden, CT) was filled with 3 M KCl and impaled an oocyte to form a tip resistance of 0.3-2 MΩ. The Geneclamp 500B amplifier and Digidata 1322A AD/DA converter (Molecular Devices, Union City, CA) were used to record the whole-cell currents and membrane potentials. The pClamp 9 software (Axon Instruments, Union City, CA) was used to acquire and analyze the data. Both signals were digitized at 200 μs/sample and filtered at 2 kHz through a Bessel filter. Data were plotted using SigmaPlot 13 (Systat Software, San Jose, CA) or GraphPad Prism 8 (GraphPad Software, San Diego, CA).

### Patch clamp

Patch-clamp recordings were carried out as we described previously (Fatehi et al., 2017), under voltage clamp at room temperature (~22°C). The recording pipette and chamber were coupled by Ag/AgCl electrodes to an Axopatch 200B patch-clamp amplifier and Digidata 1200A BNC data-acquisition system, controlled by pCLAMP 10 software (Axon Instruments). Both bath and pipette solutions were composed of (in mM) 140 NaCl, 5 KCl, 2 CaCl_2_, 2 MgCl_2_, 10 HEPES, 10 glucose and pH 7.4. The seal resistance was no less than 5 GΩ for all cell-attached recordings.

The recordings were low-pass filtered at 1 kHz following acquisition. Linear stepwise negative pressures (by suction) of various magnitudes were applied to the interior of the glass pipette using a high-speed pressure clamp system (HSPC-1, ALA Scientific, Farmingdale, NY).

### Western blot and surface protein biotinylation

*Xenopus* oocytes after 3-times washing with ice-cold PBS solution were incubated with 0.5 mg/ml sulfo-NHS-SS-Biotin (Pierce, Rockford, IL) for 30 min at room temperature. Non-reacted biotin was quenched by 1 M NH_4_Cl. Oocytes were then washed with ice-cold PBS solution and harvested in ice-cold CelLytic M lysis buffer (Sigma-Aldrich, St. Louis, MO) supplemented with proteinase inhibitor cocktail (Thermo Fisher Scientific). Upon addition of 100 μl streptavidin (Pierce) lysates were incubated at 4 °C overnight with gentle shaking. The surface protein bound to streptavidin was resuspended in SDS loading buffer and subjected to SDS-PAGE.

### Chemical cross-linking

Chemical cross-linking assays were performed as previously described (Yu et al., 2009). Oocytes expressing desired proteins were harvested in ice-cold CelLytic M lysis buffer (Sigma-Aldrich) supplemented with proteinase inhibitor cocktail (Thermo Fisher Scientific). Fresh crosslinker stock solutions were prepared by dissolving DST (CovaChem) into dimethylsulfoxide (Sigma-Aldrich). Cross-linking reactions were carried out by diluting the stock solution into the cell lysate samples, followed by incubation on ice for 4 h. SDS loading buffer was used to stop the reactions. The samples were incubated at 37 °C for 30 min and subjected to SDS-PAGE.

### Immunofluorescence

Whole-mount immunofluorescence assays using *Xenopus* oocytes were performed as described (Zheng et al., 2018a). Briefly, oocytes were washed in PBS, fixed in 4% paraformaldehyde for 15 min, washed three times in PBS plus 50 mM NH_4_Cl, and then permeabilized with 0.1% Triton X-100 for 4 min. Oocytes were then blocked in PBS plus 3% skim milk for 30 min and then incubated overnight with indicated primary antibodies, followed by incubation with secondary Alexa-488-conjugated donkey anti-rabbit or Cy3-conjugated goat anti-mouse antibody (Jackson ImmunoResearch Laboratories, West Grove, PA) for 30 min. Oocytes were then mounted in Vectashield (Vector Labs, Burlington, ON, Canada) and examined on an AIVI spinning disc confocal microscopy (Cell Imaging Facility, Faculty of Medicine and Dentistry, University of Alberta).

### Co-IP

Co-IP experiments were performed as we previously described (Zheng et al., 2018a). Briefly, a group of 20-30 oocytes washed with PBS were solubilized in ice-cold CelLytic-M lysis buffer (Sigma-Aldrich) supplemented with proteinase inhibitor cocktail. Supernatants were collected after centrifugation at 13,200 rpm for 15 min and precleaned for 1 h with 50% protein G-Sepharose (GE Healthcare), followed by incubation with an indicated antibody at 4 °C Upon addition of 100 μl of 50% protein G-Sepharose, the mixture was incubated at 4°C for overnight with gentle shaking. The immune complexes conjugated to protein G-Sepharose were washed three times with cold PBS solution containing 1% Nonidet P-40 and eluted by SDS loading buffer. Precipitated proteins were subjected to Western blot analysis.

### Zebrafish experiments

Zebrafish experiments were modified from one described previously (Zheng et al., 2018b). Briefly, embryos of WT zebrafish AB line were grown at 28.5 °C in water supplemented with 60 μg/ml Instant Ocean Sea salts. Single guide RNAs (sgRNAs: AGAACACCGCTCTAGTGCCG) targeting the first coding exon of zebrafish PKD2 was designed and synthesized. The Cas9 protein was purchased from NEB. Mixture of sgRNA (100 pg) and Cas9 (200 pg) was injected into each fertilized embryo at 1-2 hpf for PKD2 KD. The effect of injected CRISPR/Cas9 was confirmed by sequencing. Human TACAN mRNA was injected into fertilized embryos at 1-2 hpf at 100 pg each. This study has been approved by the Hubei University of Technology animal welfare regulations and maintained according to standard protocols (http://ZFIN.org).

Histological analysis was performed as previously reported (Wang et al., 2020). Briefly, embryos at 3 dpf were anesthetized in tricaine solution and fixed in 4% PFA at 4 °C. After 3-times wash (20 min each) in cold PBS, embryos were decalcified in EDTA solution (684 mM EDTA, pH 8.0) and then washed (2 × 20 min) with DEPC solution and dehydrated through graded alcohol. Before paraffin processing, embryos were embedded in 1% agarose (Sigma-Aldrich) in TAE buffer (Thermo Fisher Scientific). Subsequently, samples were sectioned transversely at 5 μm thickness using a HM325 manual rotary microtome (Thermo Fisher Scientific). H&E staining was processed with Varistain TM Gemini ES Automated Slide Stainer (Thermo Fisher Scientific).

### Statistical analysis

All statistical data in this study were represented as mean ± SEM (standard error of the mean) from N measurements. Student’s t-test was used for two groups comparisons and one-way ANOVA for multiple groups comparisons. *, ** and *** indicate p < 0.05, 0.01 and 0.001, respectively; ns indicates statistically not significant.

## Author contributions

Conceptualization, X.L., and X.-Z.C.; Investigation, X.L., R.Z., M.F., Y.W., W.L., R.T., X.D., Z.W.; Supervision, P.R.L., J.T., and X.-Z.C. Writing, X.L., and X.-Z.C.

## Acknowledgments

We would like to thank Dr. Reza Sharif-Naeini for sharing TACAN related plasmids. This work was supported by the Natural Sciences and Engineering Research Council of Canada (NSERC), the Kidney Foundation of Canada (to X.Z.C.) and National Natural Science Foundation of China (grant # 31871176 and 32070726, to J.T.). X.L. was a recipient of the Alberta Graduate Excellence Scholarship.

## Declaration of Interests

The authors declare no competing interests.

---

**Figure S1.**
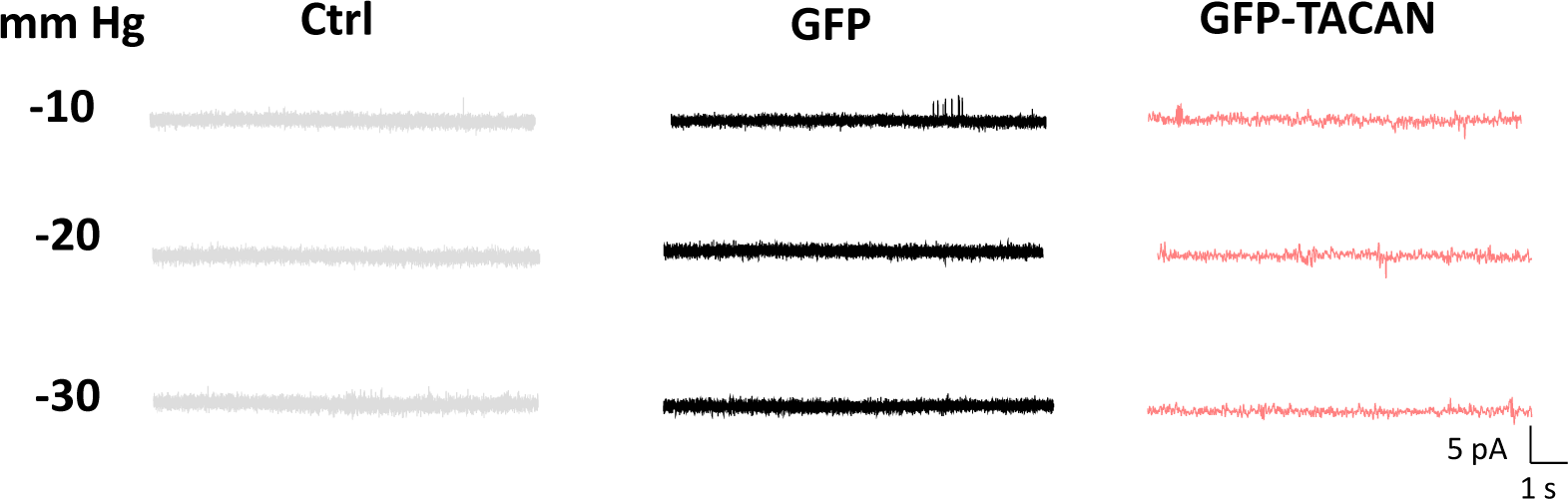
Channel activity of TACAN in CHO cells under negative pressures, related to Figure 2. Representative single channel recordings in CHO cells transfected with none (Ctrl), GFP or GFP-TACAN, in the presence of a negative pressure, as indicated.

**Figure S2.**
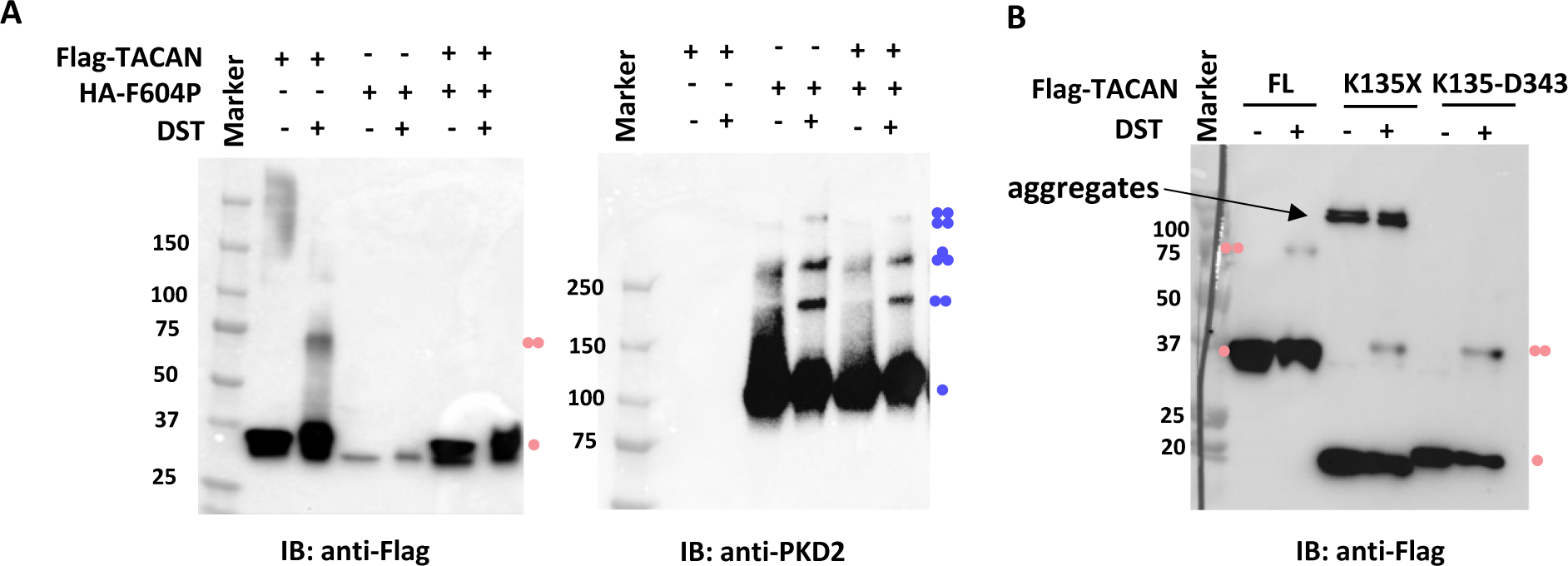
Oligomerization of TACAN and PKD2 F604P, related to Figure 5. **A.** Western blot data obtained in the presence of cross-linking, showing expression of TACAN and PKD2 F604P. Cross-linking was carried out with 1 mM disuccinimidyl tartrate (DST). Putative subunit composition of the bands is indicated. B. Western blot data obtained in the presence of cross-linking, showing expression of TACAN FL, K135X, and K135X-D343. Putative oligomer conditions are indicated.

## Notes

### Competing Interest Statement

The authors have declared no competing interest.

